# TooT-SC: Predicting Eleven Substrate Classes of Transmembrane Transport Proteins

**DOI:** 10.1101/2022.01.25.477715

**Authors:** Munira Alballa, Gregory Butler

## Abstract

**Background:** Transporters form a significant proportion of the proteome and play an important role in mediating the movement of compounds across membranes. Transport proteins are difficult to characterize experimentally, so there is a need for computational tools that predict the substrates transported in order to annotate the large number of genomes being sequenced. Recently we developed a dataset of eleven substrate classes from `Swiss-Prot` using the `ChEBI` ontology as the basis for the definition of the classes.

**Results:** We extend our earlier work *TranCEP*, which predicted seven substrate classes, to the new dataset with eleven substrate classes. Like *TranCEP, TooT-SC* combines pairwise amino acid composition (PAAC) of the protein, with evolutionary information captured in a multiple sequence alignment (MSA) using TM-Coffee, and restriction to important positions of the alignment using TCS. Our experimental results show that *TooT-SC* significantly outperforms the state-of-the-art predictors, including our earlier work, with an overall MCC of 0.82 and the MCC for the eleven classes ranging from 0.66 to 1.00.

**Conclusion:** *TooT-SC* is a useful tool with high performance covering a broad range of substrate classes. The results quantify the contribution made by each type of information used during the prediction process. We believe the methodology is applicable more generally for protein sequence analysis.

## Background

Transport proteins play important roles in biological processes [1] and form a large proportion of all proteins in an organism [2], yet existing tools for the annotation of transporters that predict the substrates of transport reactions lag behind tools for other kinds of proteins, such as for predicting enzymes involved in metabolic reactions. Many tools rely simply on homology or orthology to predict transporters. These tools include the metabolic network tools merlin [3–5], Pantograph [6], and TransATH [7] that process the complete proteome and predict each transport reaction, which means identifying the transport protein and the specific substrate, as well as cellular compartment.

Among the tools for *de novo* prediction of substrate class, FastTrans [8] claims to be the state-of-the-art. The *de novo* prediction tools predict the type of substrate from a general subset of substrate types, without attempting to predict the specific substrate [9–13], due to the limited number of transporters annotated with specific substrates. Until now, these tools have reached a maximum of seven substrate types [13] [8].

In 2019 we developed a dataset [14] that defined substrate classes in terms of the `ChEBI` ontology for Chemical Entities of Biological Interest [15]. Transporters in `Swiss-Prot` that have‘ `GO` annotations of functional transport activity of a substrate contain a link to the `ChEBI` term for the particular substrate as part of the `GO` term. The `ChEBI` hierarchy allowed us to group substrates into classes giving us eleven well-defined classes with sufficient number of examples for machine learning. To the best of our knowledge, these data contain the highest number of substrate classes being used to predict the substrate class of a transporter.

This paper extends our previous work *TranCEP* [16]. This work follows the same methodology, however, using the new dataset with eleven classes. As before we studied the impact of protein composition, protein evolution, and the specificity-determining positions within the protein sequence. The best approach, which defines *TooT-SC*, involves utilizing the PAAC encoding scheme, the TM-Coffee MSA algorithm [**?**], and the transitive consistency score (TCS) algorithm [17] to create vectors as input to build a suite of SVM classifiers, one for distinguishing each substrate class. The difference between the work on *TranCEP* and *TooT-SC* are

- the different datasets with eleven versus seven substrate classes;
- the definition of substrate classes using the `ChEBI` ontology;
- using the `Swiss-Prot` annotation of the substrate of a transport protein rather than manual annotation by the researchers;
- building the multi-class SVM classifier as a collection of one-versus-rest binary SVM classifiers (like TrSSP [13]) rather than as one-versus-one classifiers; and
- using the SVM probabilities to classify a test protein.

Readers seeking more background on work in this area, and the details of the methodology are referred to our previous paper on *TranCEP* [16] and the PhD thesis [18] of the first author.

## Materials and Methods

### Dataset

The dataset was constructed from `Swiss-Prot` using the `ChEBI` ontology [15] as described in [14]. The dataset contains 11 substrate classes, with the largest being the *inorganic cations* class with 601 samples and the smallest being the *nucleotide* class with 24 samples, as presented in Table 1. The data were randomly partitioned (stratified by class) into training (90%) and testing (10%) sets. We refer to the data in Table 1 as *DS-SC*.

**Table 1.** Dataset *DS-SC*

| ID | Substrate class | Training | Testing | Total |
| --- | --- | --- | --- | --- |
| C1 | Nonselective | 24 | 2 | 26 |
| C2 | Water | 24 | 2 | 26 |
| C3 | Inorganic cations | 541 | 60 | 601 |
| C4 | Inorganic anions | 92 | 10 | 102 |
| C5 | Organic anions | 97 | 10 | 107 |
| C6 | Organo-oxygens | 157 | 17 | 174 |
| C7 | Amino acids and derivatives | 142 | 15 | 157 |
| C8 | Other organonitrogens | 144 | 16 | 160 |
| C9 | Nucleotides | 22 | 2 | 24 |
| C10 | Organic heterocyclics | 34 | 3 | 37 |
| C11 | Miscellaneous | 99 | 11 | 110 |
| **Total** |  | **1,376** | **148** | **1,524** |

### Databases

We used the same databases as before: `Swiss-Prot` database when searching for similar sequences; and the `UniRef50-TM` database, which consists of the entries in `UniRef50` that have the keyword *transmembrane*, inside TM-Coffee [19] when constructing MSAs. Since dataset was derived from `Swiss-Prot`, we removed the exact hits of test sequences from the two databases `Swiss-Prot` and `UniRef50-TM`.

### Algorithm

Algorithm 1 presents the template for constructing the vectors required for the SVM classifiers. It combines evolutionary (E), positional (P), and compositional (C) information. The first two are optional. We used TM-Coffee to compute the MSA that conserves the TMSs and the TCS to determine a reliability index for each position (column) in the MSA. We experimented with three composition schemes, AAC, PAAC, and PseAAC, as well as the optional use of TM-Coffee and the TCS.

#### Algorithm 1

Template for constructing the composition vector

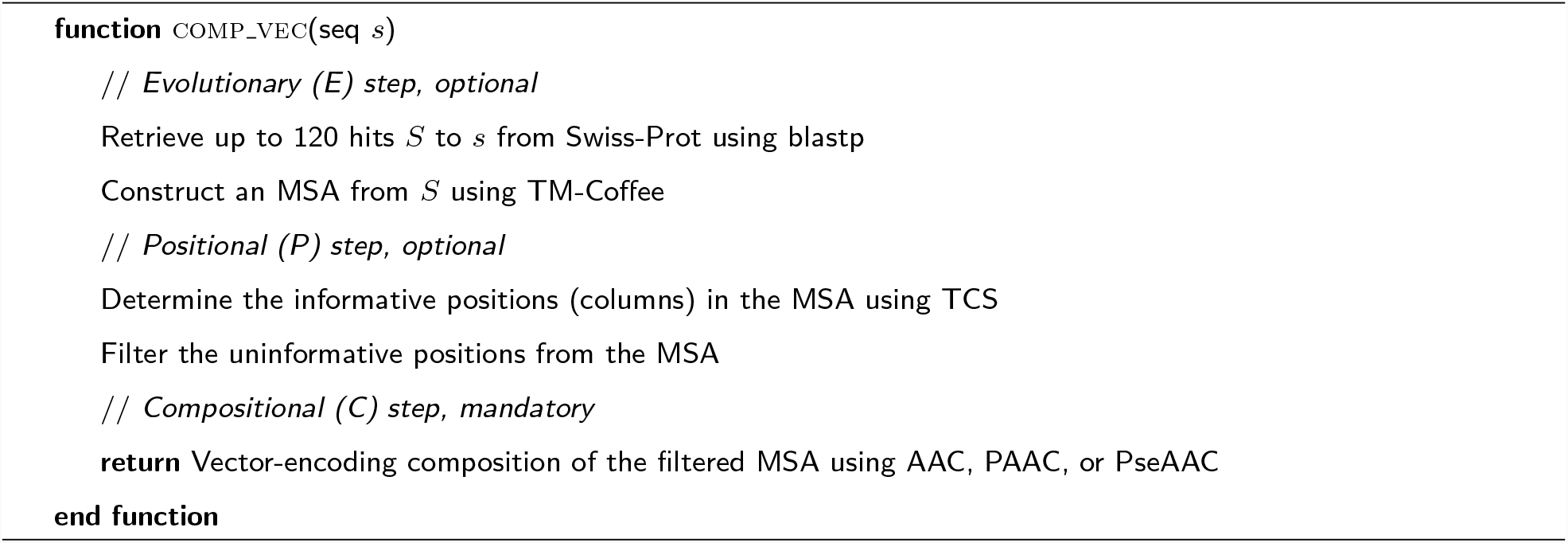

Algorithm 2 shows the composition vectors being used to build a set of SVM classifiers. In this case, multi-class classification is done using a collection of binary classifiers as one-versus-rest for each of the eleven classes.

#### Algorithm 2

Building the SVM classifiers

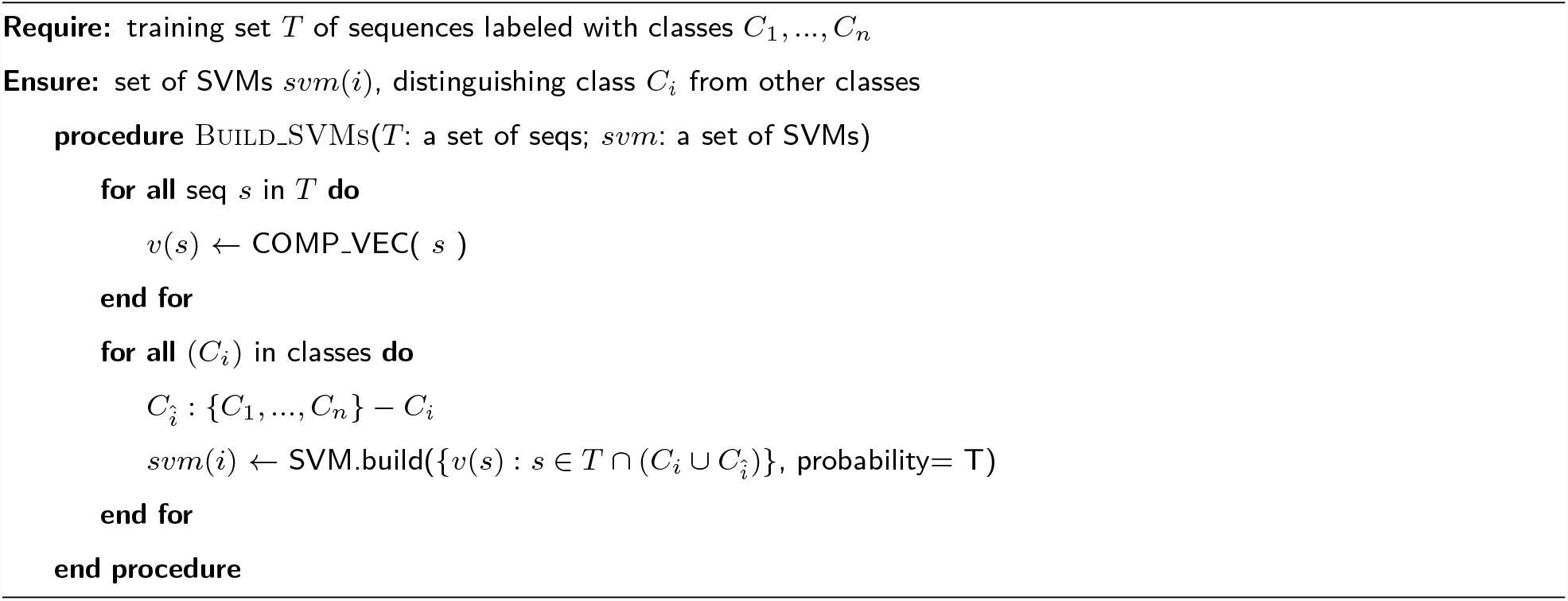

Algorithm 3 presents the prediction algorithm. Here we use the probability of each class prediction as returned by the SVM to determine the classification.

#### Algorithm 3

Prediction

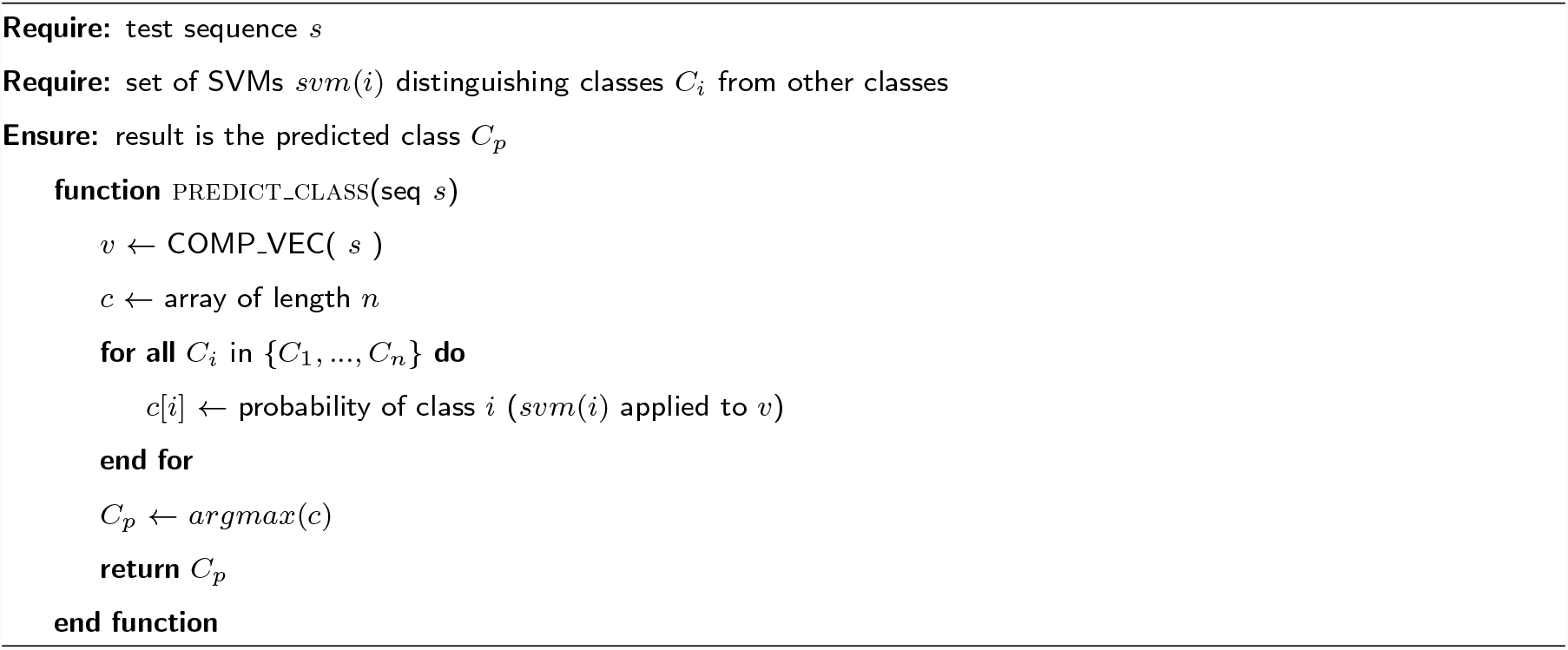

### Training

We used SVM with an RBF kernel, as implemented in the `R` *e1071* library (version 1.6-8), utilizing a one-against-the-rest approach in which *n* binary classifiers are trained, one for each class. The classifier *i* is trained with all the samples of class *i* as a positive class and the rest as a negative class. The final predicted class is the class with the highest probability among the *n* predictions. Both the cost and *γ* parameters of the RBF kernel were optimized by performing a grid search using the *tune* function in the library (cost range: 2^(1…5)^, *γ* range: 2^(−18…2)^).

### Methods

We experimented with nine methods with different combinations of inforamation:

- **AAC, PAAC**, and **PseAAC** using only compositional information;
- **TMC-AAC, TMC-PAAC**, and **TMC-PseAAC** using evolutionary and compositional information; and
- **TMC-TCS-AAC, TMC-TCS-PAAC**, and **TMC-TCS-PseAAC** using evolutionary, positional, and compositional information.

The method used in ***TooT-SC*** is **TMC-TCS-PAAC**, the method that achieved the best performance during cross-validation.

### Performance evaluation

The performance of each method on the *DS-SC* training set was determined using ten-fold cross-validation (10-CV). We repeated the 10-CV process ten times with different random partitions, to make the error estimation more stable, and reported the performance variations between the runs by computing the standard deviation.

Four performance metrics were considered:

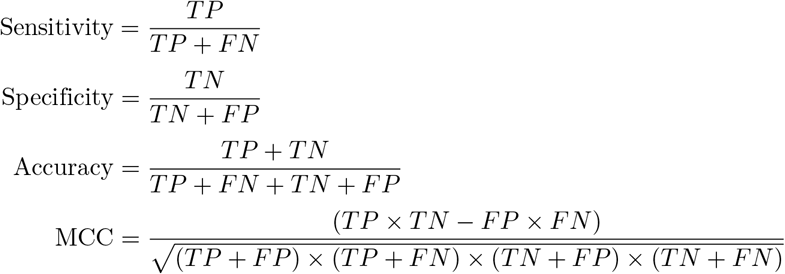

where *TP* is the number of true positives, *TN* is the number of true negatives, *FP* is the number of false positives, and *FN* is the number of false negatives.

The Matthews Correlation Coefficient (MCC) is less influenced by imbalanced data and is arguably the best single assessment metric in this case [20–22]. The overall performance across all classes was the micro-average of the individual results due to the imbalanced dataset.

### Statistical analysis

In this analysis, Student’s (two-tailed, paired) t-tests were applied, and the average number of informative residues, as determined by TCSs, in different segments of a protein sequence was computed. For each substrate class, pairwise comparisons between the means of important positions in different segments were performed. The differences were considered statistically significant when the P-value of the Student’s t-test was less than 0.0001.

## Results and Discussion

### Methods evaluation

Since the data are imbalanced, we focused on the MCC when comparing the performances of the different models. Table 2 presents the overall accuracy values and MCCs of the SVM models for the nine methods, sorted from the best to the worst according to the MCC. The details of the performance for each method are available in Supplementary Material 1. the comparisons among the different methods for the eleven classes in terms of the MCC are presented in Figure 1. The SVM model that utilized PAAC encoding outperformed those that utilized AAC and PseAAC encoding by 27% and 15%, respectively, in terms of the overall MCCs. This model shows exceptionally high performance in the *water* and *nucleotide* classes. In addition, all of the SVM models that utilized evolutionary data performed notably better overall than the SVM models that did not. The top model, **TMC-TCS-PAAC**, which is the method chosen for our predictor ***TooT-SC***, incorporates the use of the PAAC with evolutionary data in the form of MSA with positional information, in which columns that have a reliability below 4 are filtered out. We found that the performance peaked using this threshold and started to decline when columns with a reliability index greater than 4 were filtered out. The **TMC-TCS-PAAC** method yielded an overall MCC of 0.77 during cross-validation. Table 4 shows the impact of evolutionary information and positional information on the composition-encoding PAAC.

**Table 2.** Overall cross-validation performance of the methods. For each method, the table presents the accuracy and MCC as the *mean ± SD* across the ten runs of the 10-fold cross-validation.

| Method | Accuracy | MCC |
| --- | --- | --- |
| **TMC-TCS-PAAC** | 82.53 ± 0.12 | 0.7772 ± 0.0019 |
| **TMC-PAAC** | 81.92 ± 0.12 | 0.7695 ± 0.0014 |
| **TMC-AAC** | 79.84 ± 0.13 | 0.7430 ± 0.0014 |
| **TMC-PseAAC** | 79.46 ± 0.30 | 0.7374 ± 0.0038 |
| **TMC-TCS-AAC** | 79.33 ± 0.24 | 0.7360 ± 0.0035 |
| **TMC-TCS-PseAAC** | 79.03 ± 0.27 | 0.7324 ± 0.0037 |
| **PAAC** | 58.93 ± 0.45 | 0.4610 ± 0.0069 |
| **PseAAC** | 54.80 ± 0.76 | 0.3999 ± 0.0108 |
| **AAC** | 52.21 ± 0.60 | 0.3628 ± 0.0091 |

**Table 3.** Detailed *TooT-SC* performance. The table presents the performance as the *mean ± SD* across the ten runs of the 10-fold cross-validation.

| Class ID | Sensitivity | Specificity | Accuracy | MCC |
| --- | --- | --- | --- | --- |
| C1 | 75.00 ± 0.00 | 99.78 ± 0.00 | 99.21 ± 0.00 | 0.7979 ± 0.0000 |
| C2 | 95.83 ± 0.00 | 99.85 ± 0.00 | 99.74 ± 0.00 | 0.9376 ± 0.0000 |
| C3 | 95.19 ± 0.47 | 86.92 ± 0.28 | 89.36 ± 0.21 | 0.7936 ± 0.0046 |
| C4 | 64.35 ± 1.97 | 99.24 ± 0.18 | 96.38 ± 0.19 | 0.7252 ± 0.0155 |
| C5 | 68.04 ± 0.49 | 98.40 ± 0.13 | 95.66 ± 0.14 | 0.6974 ± 0.0084 |
| C6 | 83.44 ± 0.52 | 98.97 ± 0.12 | 96.72 ± 0.15 | 0.8543 ± 0.0066 |
| C7 | 84.08 ± 0.95 | 98.55 ± 0.16 | 96.56 ± 0.18 | 0.8357 ± 0.0085 |
| C8 | 71.46 ± 0.95 | 96.84 ± 0.27 | 93.42 ± 0.22 | 0.6830 ± 0.0084 |
| C9 | 80.91 ± 1.92 | 99.98 ± 0.04 | 99.61 ± 0.05 | 0.8904 ± 0.0132 |
| C10 | 82.35 ± 0.00 | 100.00 ± 0.00 | 99.47 ± 0.00 | 0.9050 ± 0.0000 |
| C11 | 55.96 ± 1.09 | 97.95 ± 0.16 | 94.21 ± 0.20 | 0.5858 ± 0.0136 |
| **Overall** | | | 82.53 ± 0.12 | 0.7772 ± 0.0019 |

**Fig 1.**
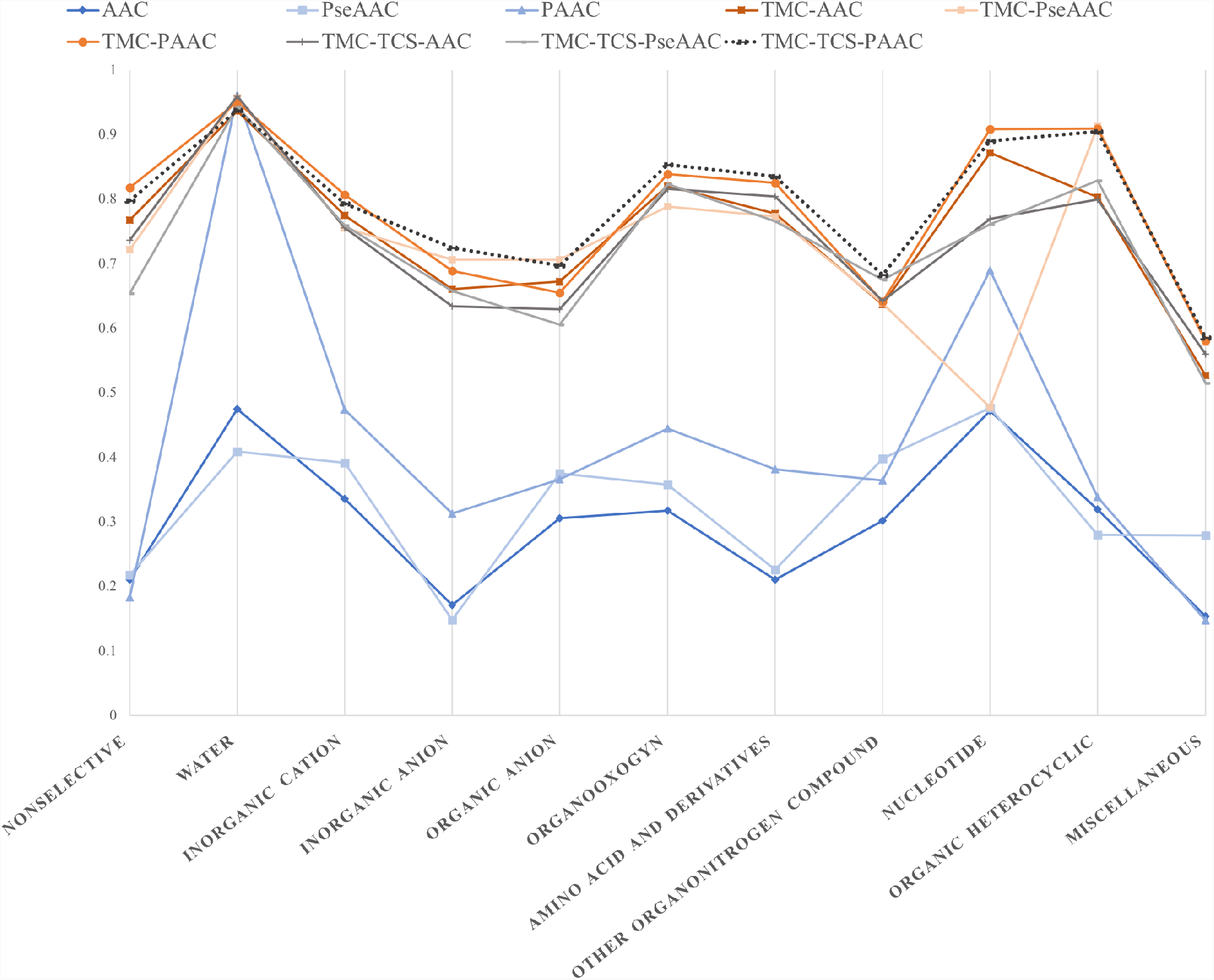
Performance of methods on the substrate classes. This figure shows the cross-validation MCC performance of the different methods on the eleven substrate classes. The dotted line represents the performance of *TooT-SC*, which is TCS-TMC-PAAC.

**Table 4.**
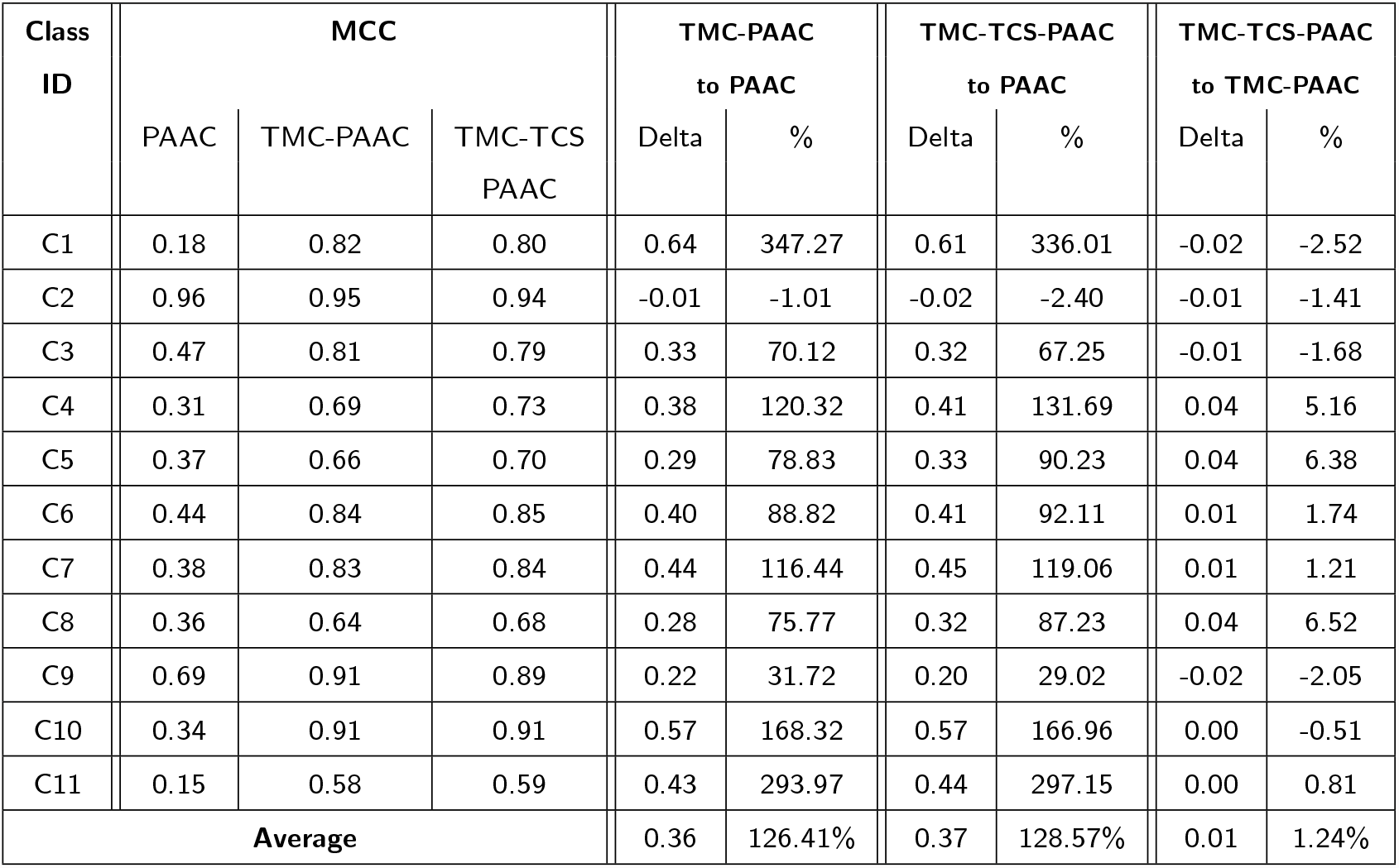
Impact of factors on performance for PAAC. This table notes the MCC and the differences in the MCC for the cross-validation performance of the methods using evolutionary information with TM-Coffee, and positional information with TCS. The differences in MCC are shown in the Delta column. The percentage improvement (loss) is also shown. The use of evolutionary information in the form of an MSA on the composition-encoding **PAAC** improved the MCC by an average of 126.41%. The further use of positional information by filtering out the unreliable columns from the MSA boosted the MCC of the composition encodings by an average of 128.57%.

| Class ID | MCC |  |  | TMC-PAAC to PAAC |  | TMC-TCS-PAAC to PAAC |  | TMC-TCS-PAAC to TMC-PAAC |  |
| --- | --- | --- | --- | --- | --- | --- | --- | --- | --- |
|  | PAAC | TMC-PAAC | TMC-TCS PAAC | Delta | % | Delta | % | Delta | % |
| C1 | 0.18 | 0.82 | 0.80 | 0.64 | 347.27 | 0.61 | 336.01 | -0.02 | -2.52 |
| C2 | 0.96 | 0.95 | 0.94 | -0.01 | -1.01 | -0.02 | -2.40 | -0.01 | -1.41 |
| C3 | 0.47 | 0.81 | 0.79 | 0.33 | 70.12 | 0.32 | 67.25 | -0.01 | -1.68 |
| C4 | 0.31 | 0.69 | 0.73 | 0.38 | 120.32 | 0.41 | 131.69 | 0.04 | 5.16 |
| C5 | 0.37 | 0.66 | 0.70 | 0.29 | 78.83 | 0.33 | 90.23 | 0.04 | 6.38 |
| C6 | 0.44 | 0.84 | 0.85 | 0.40 | 88.82 | 0.41 | 92.11 | 0.01 | 1.74 |
| C7 | 0.38 | 0.83 | 0.84 | 0.44 | 116.44 | 0.45 | 119.06 | 0.01 | 1.21 |
| C8 | 0.36 | 0.64 | 0.68 | 0.28 | 75.77 | 0.32 | 87.23 | 0.04 | 6.52 |
| C9 | 0.69 | 0.91 | 0.89 | 0.22 | 31.72 | 0.20 | 29.02 | -0.02 | -2.05 |
| C10 | 0.34 | 0.91 | 0.91 | 0.57 | 168.32 | 0.57 | 166.96 | 0.00 | -0.51 |
| C11 | 0.15 | 0.58 | 0.59 | 0.43 | 293.97 | 0.44 | 297.15 | 0.00 | 0.81 |
| **Average** |  |  |  | 0.36 | 126.41% | 0.37 | 128.57% | 0.01 | 1.24% |

The use of evolutionary information in the form of MSA on the composition-encoding PAAC showed a considerable positive impact in most of the substrate classes, where the average improvement of the MCC was 126.41%, with the highest improvement being in the C1 (*nonselective*) class (347%). The baseline encoding PAAC for the C2 (*water*) substrate class showed a high discriminatory power with an MCC of 0.96, with the incorporation of additional information having a slightly negative impact of 1.01%.

The further use of positional information by filtering out the unreliable columns from the MSA showed an average improvement of 128.57% compared to the baseline compositions. The impact of positional information over that already achieved by evolutionary information showed a positive impact in most substrate classes; the highest was in the C5 (*organic anions*) class, where the MCC improved by 6.38% with **TMC-TCS-PAAC**. However, the impact was slightly negative in the C1 (*nonselective*), C2 (*water*), C3 (*inorganic cations*), and C9 (*nucleotides*) classes.

### Comparison with other published work

The top two tools with the best reported performance are TrSSP [13] and FastTrans [8]. Since the original code was not available for TrSSP or FastTrans, we reimplemented the methods to the best of our ability. We compared the performance of the *TooT-SC* method with our implementation of the TrSSP and FastTrans methods. All of the methods were trained using the *DS-SC* training set and tested using its testing set. It should be noted that our implementation of the TrSSP method [13] achieved a similar macroaverage MCC to that reported in the original paper (0.41) on their dataset. However, it was not possible to reproduce the reported performance of the FastTrans method [8], for which our implementation on their same dataset achieved a macroaverage MCC of 0.47, while their reported macroaverage MCC was 0.87.

A comparison between the *TooT-SC* method and our implementation of the other state-of-the-art methods on the *DS-SC* benchmark dataset is presented in Table 5. The *TooT-SC* method scored higher than the other methods for all of the substrate classes in terms of the accuracy, sensitivity, and MCC. Overall, the *TooT-SC* method scored an overall MCC of 0.82, which outperformed the TrSSP method by 26% and the FastTrans method by 115%.

**Table 5.** Comparison between *TooT-SC* and the state-of-art methods. This table presents the performance of the proposed tool *TooT-SC* built with the complete training set and run on the independent testing set of *DS-SC* and the corresponding results for the TrSSP and FastTrans methods trained and tested with the same dataset. This table shows the specificity, sensitivity, accuracy and MCC for each of the eleven substrate types; the overall accuracy and MCC; and the macroaverage accuracy and MCC. The overall accuracy was calculated as the number of correct predictions divided by the total number of predictions, and the overall MCC was calculated from the multi-class confusion matrix.

| Class ID | Specificity |  |  | Sensitivity |  |  | Accuracy |  |  | MCC |  |  |
| --- | --- | --- | --- | --- | --- | --- | --- | --- | --- | --- | --- | --- |
|  | TrSSP | FastTrans | TooT-SC | TrSSP | FastTrans | TooT-SC | TrSSP | FastTrans | TooT-SC | TrSSP | FastTrans | TooT-SC |
| C1 | 100.00 | 100.00 | 100.00 | 0.00 | 0.00 | 50.00 | 98.18 | 97.70 | 99.22 | 0.00 | 0.00 | 0.70 |
| C2 | 99.32 | 100.00 | 100.00 | 100.00 | 100.00 | 100.00 | 99.08 | 100.00 | 100.00 | 0.81 | 1.00 | 1.00 |
| C3 | 80.68 | 76.14 | 88.64 | 91.67 | 86.67 | 96.67 | 83.08 | 74.56 | 91.37 | 0.68 | 0.50 | 0.83 |
| C4 | 98.55 | 95.65 | 100.00 | 60.00 | 40.00 | 70.00 | 94.74 | 87.63 | 97.69 | 0.64 | 0.33 | 0.83 |
| C5 | 98.55 | 97.83 | 97.83 | 80.00 | 50.00 | 90.00 | 96.43 | 91.40 | 96.95 | 0.78 | 0.51 | 0.81 |
| C6 | 96.95 | 96.18 | 97.71 | 64.71 | 35.29 | 76.47 | 91.53 | 84.16 | 94.78 | 0.64 | 0.35 | 0.76 |
| C7 | 97.74 | 87.97 | 100.00 | 73.33 | 40.00 | 86.67 | 93.91 | 77.27 | 98.45 | 0.72 | 0.20 | 0.92 |
| C8 | 94.70 | 96.21 | 96.21 | 56.25 | 25.00 | 87.50 | 88.52 | 83.33 | 94.78 | 0.50 | 0.25 | 0.77 |
| C9 | 99.32 | 99.32 | 100.00 | 100.00 | 0.00 | 100.00 | 99.08 | 96.59 | 100.00 | 0.81 | -0.02 | 1.00 |
| C10 | 98.62 | 100.00 | 100.00 | 33.33 | 66.67 | 100.00 | 96.43 | 98.84 | 100.00 | 0.31 | 0.81 | 1.00 |
| C11 | 99.27 | 95.62 | 100.00 | 27.27 | 36.36 | 45.45 | 92.31 | 86.73 | 95.49 | 0.42 | 0.31 | 0.66 |
| **Overall** |  |  |  |  |  |  | 72.97 | 57.43 | 85.81 | 0.65 | 0.44 | 0.82 |
| **Macroaverage** |  |  |  |  |  |  | 93.94 | 88.93 | 97.16 | 0.57 | 0.39 | 0.84 |

### Positional information analysis

See Supplementary Material 2.

## Conclusion

We have developed *TooT-SC* for the *de novo* prediction of substrates for membrane transporter proteins that combines information based on the amino acid composition, evolutionary information, and positional information. *TooT-SC* is able to classify transport proteins into eleven classes according to their transported substrate (i.e., *nonselective, water, inorganic cations, inorganic anions, organic anions, organo-oxygens, amino acids and derivatives, other organonitrogens, nucleotides, organic heterocyclics*, and *miscellaneous*); to the best of our knowledge, this is the highest number of classes offered by a *de novo* prediction tool. The *TooT-SC* method first incorporates the use of evolutionary information by taking 120 similar sequences and constructing an MSA using TM-Coffee. Next, it uses the positional information by filtering out unreliable positions, as determined by the TCS, and then uses the PAAC. The *TooT-SC* method achieved an overall MCC of 0.82 on an independent testing set, which is a 26% improvement over the state-of-the-art method. In addition, we evaluated the impact of each factor on the performance by incorporating evolutionary information and filtering out unreliable positions. We observed that the PAAC encoding outperforms other combinational variations. However, it does not show compelling performance on its own; the enhanced performance comes mainly from incorporating evolutionary and positional information.

Analysis of the location of the informative positions reveals that there are more statistically significant informative positions in the TMSs compared to the non-TMSs and there are more statistically significant informative positions that occur close to the TMSs compared to regions far from them.

In moving from the previous gold standard dataset with seven substrate classes to our new dataset with eleven substrate classes, even with the same approach, the overall MCC rose from 0.69 to 0.82. The impact of the positional information is statistically significant more often with the new dataset. The datasets do use different classes, however, we would like to think that the improvement is due to using substrate classes defined in terms of the `ChEBI` ontology, and selecting proteins in `Swiss-Prot` with curated `GO` annotations clearly indicating the substrate in terms of `ChEBI`.

## Availability of data and materials

Data and materials are avialable at https://github.com/bioinformatics-group/TooT-SC

## Author’s contributions

MA developed *TranCEP, TooT-SC*, and the DS-SC dataset. MA performed the experiments, and drafted the manuscript as part of her PhD thesis under GB. GB developed the roadmap for the methodology, conceived the project, and supervised the work. All authors reviewed and approved the article.

## Acknowledgements

MA was supported by King Saud University in Riyadh, Saudi Arabia, and the Saudi Arabian Cultural Bureau in Canada. GB was partially supported by the NSERC Discovery Grant programme, and by Genome Canada, Genome Québec, and Concordia University for the Bioinformatics and Computational Biology Competition 2017.

## Additional Files

Additional file 1 — Supplementary Material 1

Tables with detailed performance of each of the nine methods on the eleven classes.

Additional file 2 — Supplementary Material 2

Results and discussion on the positional information.

## Supplementary Material 1: Detailed Performance per Class per Method

This contains the detailed cross-validation performance in substrate specificity prediction. The following tables show the *mean* ± *SD* of the ten different runs of the ten-fold cross validation for each combination of approaches.

**Table 1.** AAC Performance

| Substrate class | Sensitivity | Specificity | Accuracy | MCC |
| --- | --- | --- | --- | --- |
| Nonselective | 13.75 ± 3.95 | 99.59 ± 0.05 | 96.48 ± 0.20 | 0.2110 ± 0.0581 |
| Water | 37.92 ± 3.65 | 99.60 ± 0.06 | 97.25 ± 0.16 | 0.4748 ± 0.0372 |
| Inorganic cations | 87.08 ± 0.56 | 61.26 ± 0.88 | 64.61 ± 0.65 | 0.3364 ± 0.0124 |
| Inorganic anions | 13.80 ± 2.05 | 98.32 ± 0.31 | 87.68 ± 0.53 | 0.1716 ± 0.0349 |
| Organic anions | 32.06 ± 1.57 | 96.90 ± 0.22 | 87.18 ± 0.38 | 0.3060 ± 0.0176 |
| Organo-oxygens | 44.71 ± 2.91 | 92.42 ± 0.38 | 80.03 ± 0.61 | 0.3179 ± 0.0243 |
| Amino acids and derivatives | 30.99 ± 2.68 | 93.32 ± 0.50 | 79.93 ± 0.79 | 0.2109 ± 0.0281 |
| Other organonitrogens | 33.89 ± 1.79 | 95.56 ± 0.20 | 82.73 ± 0.36 | 0.3022 ± 0.0162 |
| Nucleotides | 41.36 ± 3.98 | 99.49 ± 0.10 | 97.32 ± 0.23 | 0.4724 ± 0.0397 |
| Organic heterocyclics | 23.82 ± 4.26 | 99.37 ± 0.11 | 95.43 ± 0.25 | 0.3194 ± 0.0440 |
| Miscellaneous | 10.91 ± 2.01 | 98.66 ± 0.24 | 87.22 ± 0.42 | 0.1539 ± 0.0370 |
| **Overall** | | | 52.21 ± 0.60 | 0.3628 ± 0.0091 |

**Table 2.**
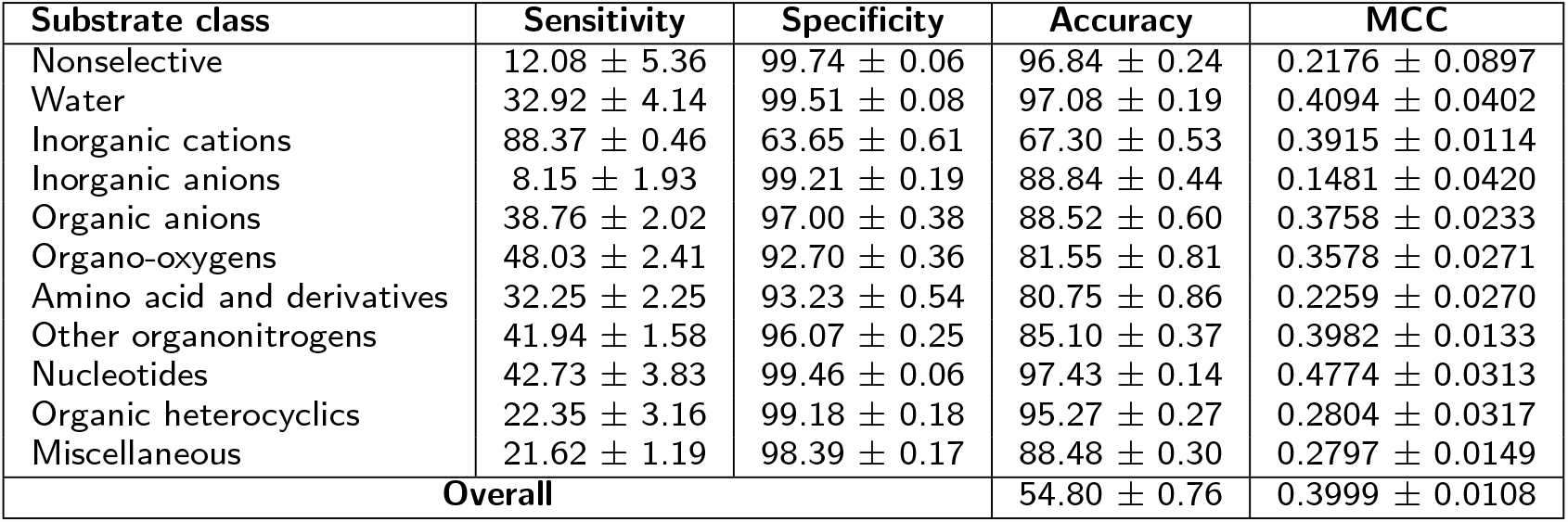
PseAAC Performance

**Table 3.** PAAC Performance

| Substrate class | Sensitivity | Specificity | Accuracy | MCC |
| --- | --- | --- | --- | --- |
| Nonselective | 10.00 ± 3.51 | 99.71 ± 0.09 | 96.95 ± 0.16 | 0.1830 ± 0.0587 |
| Water | 93.33 ± 2.91 | 99.99 ± 0.03 | 99.78 ± 0.10 | 0.9607 ± 0.0174 |
| Inorganic cations | 88.98 ± 0.65 | 69.21 ± 0.96 | 71.91 ± 0.56 | 0.4745 ± 0.0096 |
| Inorganic anions | 24.24 ± 2.35 | 98.46 ± 0.26 | 90.06 ± 0.43 | 0.3130 ± 0.0316 |
| Organic anions | 38.76 ± 1.77 | 96.69 ± 0.40 | 88.86 ± 0.49 | 0.3666 ± 0.0193 |
| Organo-oxygens | 54.20 ± 1.51 | 93.83 ± 0.51 | 84.65 ± 0.67 | 0.4447 ± 0.0211 |
| Amino acids and derivatives | 47.39 ± 2.25 | 93.91 ± 0.26 | 84.40 ± 0.43 | 0.3815 ± 0.0193 |
| Other organonitrogens | 39.58 ± 2.68 | 95.54 ± 0.33 | 85.11 ± 0.36 | 0.3648 ± 0.0194 |
| Nucleotides | 68.64 ± 3.98 | 99.54 ± 0.10 | 98.41 ± 0.21 | 0.6901 ± 0.0371 |
| Organic heterocyclics | 27.35 ± 3.41 | 99.23 ± 0.08 | 95.85 ± 0.13 | 0.3390 ± 0.0297 |
| Miscellaneous | 11.01 ± 1.93 | 98.43 ± 0.20 | 88.24 ± 0.40 | 0.1475 ± 0.0358 |
| **Overall** | | | 58.93 ± 0.45 | 0.461 ± 0.0069 |

**Table 4.**
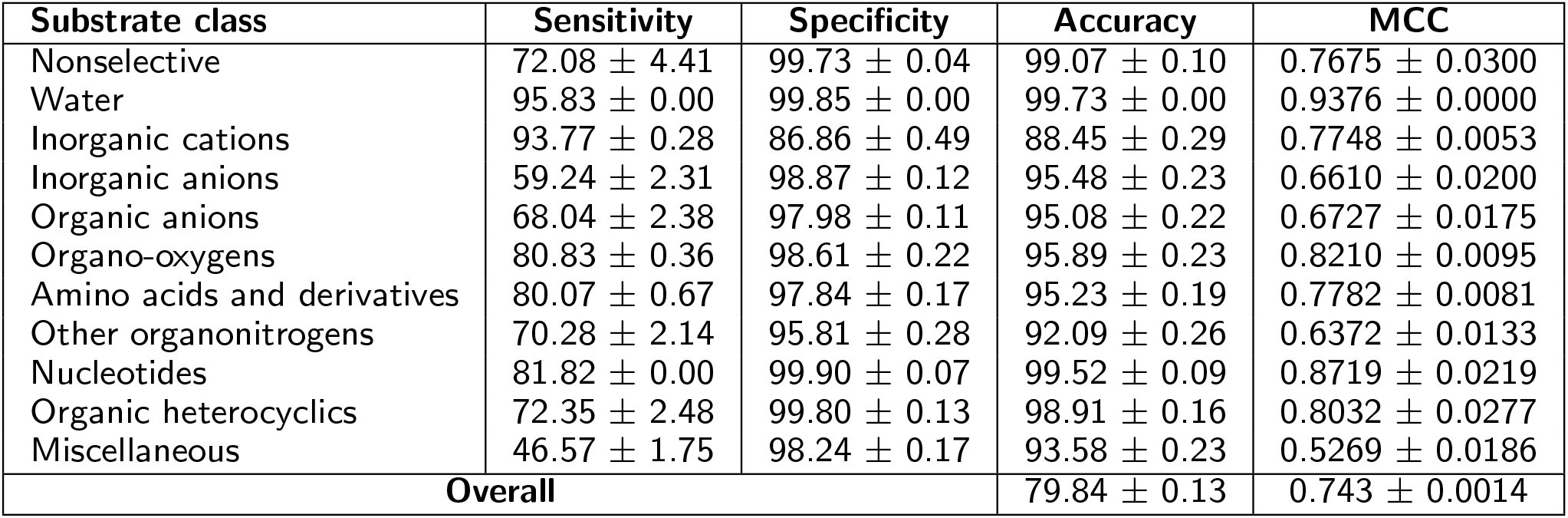
TMC-AAC Performance

| Substrate class | Sensitivity | Specificity | Accuracy | MCC |
| --- | --- | --- | --- | --- |
| Nonselective | 72.08 ± 4.41 | 99.73 ± 0.04 | 99.07 ± 0.10 | 0.7675 ± 0.0300 |
| Water | 95.83 ± 0.00 | 99.85 ± 0.00 | 99.73 ± 0.00 | 0.9376 ± 0.0000 |
| Inorganic cations | 93.77 ± 0.28 | 86.86 ± 0.49 | 88.45 ± 0.29 | 0.7748 ± 0.0053 |
| Inorganic anions | 59.24 ± 2.31 | 98.87 ± 0.12 | 95.48 ± 0.23 | 0.6610 ± 0.0200 |
| Organic anions | 68.04 ± 2.38 | 97.98 ± 0.11 | 95.08 ± 0.22 | 0.6727 ± 0.0175 |
| Organo-oxygens | 80.83 ± 0.36 | 98.61 ± 0.22 | 95.89 ± 0.23 | 0.8210 ± 0.0095 |
| Amino acids and derivatives | 80.07 ± 0.67 | 97.84 ± 0.17 | 95.23 ± 0.19 | 0.7782 ± 0.0081 |
| Other organonitrogens | 70.28 ± 2.14 | 95.81 ± 0.28 | 92.09 ± 0.26 | 0.6372 ± 0.0133 |
| Nucleotides | 81.82 ± 0.00 | 99.90 ± 0.07 | 99.52 ± 0.09 | 0.8719 ± 0.0219 |
| Organic heterocyclics | 72.35 ± 2.48 | 99.80 ± 0.13 | 98.91 ± 0.16 | 0.8032 ± 0.0277 |
| Miscellaneous | 46.57 ± 1.75 | 98.24 ± 0.17 | 93.58 ± 0.23 | 0.5269 ± 0.0186 |
| **Overall** | | | 79.84 ± 0.13 | 0.743 ± 0.0014 |

**Table 5.** TMC-PseAAC Performance

| Substrate class | Sensitivity | Specificity | Accuracy | MCC |
| --- | --- | --- | --- | --- |
| Nonselective | 63.33 ± 5.12 | 99.78 ± 0.06 | 98.93 ± 0.14 | 0.7220 ± 0.0411 |
| Water | 95.83 ± 0.00 | 99.93 ± 0.00 | 99.82 ± 0.00 | 0.9574 ± 0.0000 |
| Inorganic cations | 94.49 ± 0.59 | 84.49 ± 0.49 | 87.28 ± 0.36 | 0.7561 ± 0.0072 |
| Inorganic anions | 59.57 ± 1.76 | 99.43 ± 0.09 | 96.09 ± 0.16 | 0.7065 ± 0.0143 |
| Organic anions | 64.54 ± 1.55 | 98.94 ± 0.11 | 95.80 ± 0.14 | 0.7070 ± 0.0112 |
| Organo-oxygens | 80.19 ± 0.82 | 97.92 ± 0.23 | 95.09 ± 0.21 | 0.7887 ± 0.0081 |
| Amino acids and derivatives | 74.65 ± 1.29 | 98.55 ± 0.18 | 95.30 ± 0.23 | 0.7732 ± 0.0110 |
| Other organonitrogens | 72.50 ± 1.40 | 95.37 ± 0.48 | 91.88 ± 0.55 | 0.6390 ± 0.0207 |
| Nucleotides | 32.73 ± 4.18 | 99.79 ± 0.10 | 98.41 ± 0.12 | 0.4780 ± 0.0418 |
| Organic heterocyclics | 88.82 ± 4.11 | 99.87 ± 0.08 | 99.49 ± 0.15 | 0.9132 ± 0.0261 |
| Miscellaneous | 53.43 ± 1.30 | 98.18 ± 0.19 | 94.03 ± 0.25 | 0.5781 ± 0.0164 |
| **Overall** | | | 79.46 ± 0.30 | 0.7374 ± 0.0038 |

**Table 6.** TMC-PAAC Performance

| Substrate class | Sensitivity | Specificity | Accuracy | MCC |
| --- | --- | --- | --- | --- |
| Nonselective | 75.00 ± 0.00 | 99.85 ± 0.05 | 99.30 ± 0.06 | 0.8185 ± 0.0141 |
| Water | 98.33 ± 2.15 | 99.85 ± 0.00 | 99.79 ± 0.05 | 0.9510 ± 0.0115 |
| Inorganic cations | 95.25 ± 0.28 | 88.26 ± 0.19 | 90.11 ± 0.14 | 0.8072 ± 0.0029 |
| Inorganic anions | 63.80 ± 1.99 | 98.80 ± 0.17 | 95.86 ± 0.23 | 0.6896 ± 0.0178 |
| Organic anions | 68.04 ± 1.75 | 97.65 ± 0.06 | 94.86 ± 0.18 | 0.6556 ± 0.0146 |
| Organo-oxygens | 83.63 ± 0.67 | 98.61 ± 0.18 | 96.35 ± 0.19 | 0.8397 ± 0.0079 |
| Amino acids and derivatives | 82.96 ± 1.28 | 98.49 ± 0.10 | 96.34 ± 0.16 | 0.8257 ± 0.0085 |
| Other organonitrogens | 66.39 ± 1.68 | 96.70 ± 0.27 | 92.67 ± 0.17 | 0.6412 ± 0.0084 |
| Nucleotides | 85.45 ± 1.92 | 99.96 ± 0.06 | 99.66 ± 0.07 | 0.9090 ± 0.0185 |
| Organic heterocyclics | 83.24 ± 4.81 | 100.00 ± 0.00 | 99.50 ± 0.14 | 0.9096 ± 0.0272 |
| Miscellaneous | 54.34 ± 1.33 | 98.09 ± 0.16 | 94.18 ± 0.16 | 0.5811 ± 0.0110 |
| **Overall** | | | 81.92 ± 0.12 | 0.7695 ± 0.0014 |

**Table 7.** TMC-TCS-AAC Performance

| Substrate class | Sensitivity | Specificity | Accuracy | MCC |
| --- | --- | --- | --- | --- |
| Nonselective | 70.00 ± 2.64 | 99.66 ± 0.09 | 98.93 ± 0.13 | 0.7365 ± 0.0288 |
| Water | 100.00 ± 0.00 | 99.85 ± 0.00 | 99.82 ± 0.00 | 0.9599 ± 0.0000 |
| Inorganic cations | 92.88 ± 0.34 | 85.96 ± 0.70 | 87.52 ± 0.40 | 0.7562 ± 0.0071 |
| Inorganic anions | 54.57 ± 1.76 | 98.98 ± 0.23 | 95.21 ± 0.35 | 0.6346 ± 0.0278 |
| Organic anions | 64.43 ± 2.89 | 97.65 ± 0.17 | 94.42 ± 0.33 | 0.6294 ± 0.0245 |
| Organo-oxygens | 83.63 ± 0.31 | 98.06 ± 0.27 | 95.67 ± 0.26 | 0.8168 ± 0.0097 |
| Amino acids and derivatives | 82.68 ± 1.34 | 98.07 ± 0.18 | 95.75 ± 0.32 | 0.8049 ± 0.0149 |
| Other organonitrogens | 68.54 ± 0.98 | 96.35 ± 0.31 | 92.36 ± 0.29 | 0.6428 ± 0.0115 |
| Nucleotide | 70.45 ± 3.21 | 99.79 ± 0.11 | 99.16 ± 0.17 | 0.7698 ± 0.0415 |
| Organic heterocyclics | 70.00 ± 2.32 | 99.86 ± 0.04 | 98.90 ± 0.09 | 0.8000 ± 0.0178 |
| Miscellaneous | 49.39 ± 1.99 | 98.41 ± 0.17 | 93.94 ± 0.29 | 0.5602 ± 0.0229 |
| **Overall** | | | 79.33 ± 0.24 | 0.736 ± 0.0035 |

**Table 8.** TMC-TCS-PseAAC Performance

| Substrate class | Sensitivity | Specificity | Accuracy | MCC |
| --- | --- | --- | --- | --- |
| Nonselective | 58.33 ± 0.00 | 99.65 ± 0.04 | 98.67 ± 0.04 | 0.6545 ± 0.0089 |
| Water | 95.83 ± 0.00 | 99.87 ± 0.04 | 99.75 ± 0.04 | 0.9435 ± 0.0096 |
| Inorganic cations | 93.84 ± 0.29 | 85.50 ± 0.48 | 87.57 ± 0.33 | 0.7594 ± 0.0061 |
| Inorganic anions | 58.59 ± 1.49 | 98.90 ± 0.15 | 95.42 ± 0.13 | 0.6585 ± 0.0092 |
| Organic anions | 62.58 ± 1.46 | 97.46 ± 0.19 | 94.05 ± 0.29 | 0.6061 ± 0.0183 |
| Organo-oxygen s | 80.89 ± 0.79 | 98.67 ± 0.12 | 95.92 ± 0.11 | 0.8239 ± 0.0047 |
| Amino acids and derivatives | 78.24 ± 1.35 | 97.84 ± 0.12 | 94.98 ± 0.18 | 0.7660 ± 0.0095 |
| Other organonitrogens | 68.96 ± 1.39 | 97.15 ± 0.20 | 93.16 ± 0.22 | 0.6752 ± 0.0108 |
| Nucleotides | 76.82 ± 2.58 | 99.62 ± 0.06 | 99.06 ± 0.10 | 0.7618 ± 0.0232 |
| Organic heterocyclics | 75.00 ± 2.08 | 99.85 ± 0.05 | 99.04 ± 0.08 | 0.8294 ± 0.0145 |
| Miscellaneous | 48.89 ± 2.44 | 97.71 ± 0.18 | 93.16 ± 0.33 | 0.5158 ± 0.0255 |
| **Overall** | | | 79.03 ± 0.27 | 0.7324 ± 0.0037 |

**Table 9.** TMC-TCS-PAAC Performance

| Substrate class | Sensitivity | Specificity | Accuracy | MCC |
| --- | --- | --- | --- | --- |
| Nonselective | 75.00 ± 0.00 | 99.78 ± 0.00 | 99.21 ± 0.00 | 0.7979 ± 0.0000 |
| Water | 95.83 ± 0.00 | 99.85 ± 0.00 | 99.74 ± 0.00 | 0.9376 ± 0.0000 |
| Inorganic cations | 95.19 ± 0.47 | 86.92 ± 0.28 | 89.36 ± 0.21 | 0.7936 ± 0.0046 |
| Inorganic anions | 64.35 ± 1.97 | 99.24 ± 0.18 | 96.38 ± 0.19 | 0.7252 ± 0.0155 |
| Organic anions | 68.04 ± 0.49 | 98.40 ± 0.13 | 95.66 ± 0.14 | 0.6974 ± 0.0084 |
| Organo-oxygens | 83.44 ± 0.52 | 98.97 ± 0.12 | 96.72 ± 0.15 | 0.8543 ± 0.0066 |
| Amino acids and derivatives | 84.08 ± 0.95 | 98.55 ± 0.16 | 96.56 ± 0.18 | 0.8357 ± 0.0085 |
| Other organonitrogens | 71.46 ± 0.95 | 96.84 ± 0.27 | 93.42 ± 0.22 | 0.6830 ± 0.0084 |
| Nucleotides | 80.91 ± 1.92 | 99.98 ± 0.04 | 99.61 ± 0.05 | 0.8904 ± 0.0132 |
| Organic heterocyclics | 82.35 ± 0.00 | 100.00 ± 0.00 | 99.47 ± 0.00 | 0.9050 ± 0.0000 |
| Miscellaneous | 55.96 ± 1.09 | 97.95 ± 0.16 | 94.21 ± 0.20 | 0.5858 ± 0.0136 |
| **Overall** | | | 82.53 ± 0.12 | 0.7772 ± 0.0019 |

## Supplementary Material 2: Positional Information Analysis

It is difficult to isolate the exact residues that are key to inferring the substrate class; the results suggest that evolutionary information, obtained by MSA, is the main source for achieving a high prediction performance. In addition, the TCS informative positions (with TCSs ≥ 4) can help to filter out unnecessary noise and obtain a clearer signal to further improve the prediction. Using the TCS informative positions filtered out an average of 31% ± 19% of the sequence. However, when we attempted to filter out more positions (by using a TCS score cutoff stricter than 4), the performance started to deteriorate.

To visualize the informative positions relative to the hydropathy scale of amino acids, the hydropathy scale proposed by [1] was utilized, and the average hydropathy of each column in the MSA was computed. Higher positive scores indicate that amino acids in that region have hydrophobic properties and are likely located in a transmembrane *α*-helix segment. The TCS of each column in the alignment is noted on the hydropathy plot through color coding. Figure 1 shows diverse examples. The red shades correspond to the informative columns (TCS ≥ 4), while the gray and white shades correspond to noninformative columns that are filtered out by *TooT-SC*. In Figure 1 (a) and (b), the regions with high positive average hydropathy values appear to be more informative than those with lower values. However, in Figure 1 (c) and (d), the difference between the informative positions with high and low hydropathy values is not as clear.

To measure the informative positions relative to different segments of the protein sequence, we divided the protein sequence positions into those in the TMS and those not in the TMS. Those in the TMS were divided into the interior one-third positions, and the remaining exterior positions in the TMS. The non-TMS positions were divided into those near a TMS, that is, within 10 positions, and the remaining positions were considered far from a TMS. The location of the TMS was retrieved from the `Swiss-Prot` database under the subcellular location topology section. Table 1 shows a breakdown of where the informative positions, as determined by the TCS, are located with respect to the TMS regions.

For instance, in Figure 1 (a), 41.04% of the residues of the sequence with `UniProt`-ID `Q59NP1` are informative (i.e., correspond to informative columns in the alignment); thus, 58.96% of this sequence is filtered out. In this case, the residues in the TMSs of this protein are indeed more informative than those of the other proteins, where 100% of them are informative. On the other hand, only 29.19% of the residues in non-TMSs are informative. The difference is not as significant in the sequence with `UniProt`-ID `Q9NY37` in Figure 1 (c), where the informative positions in the TMSs are similar to those of non-TMS positions. Details of the sequences in the figure are presented in Table 2.

Table 3 presents a pairwise comparison between informative positions in the TMS and non-TMS regions. The sequences in all of the substrate classes except the C1 (*nonselective*) substrate class have significantly more informative positions in the TMS regions than in the non-TMS regions. Similarly, there is a significant difference between the informative positions close to TMSs and positions far from TMSs in all sequences that belong to all substrate classes except the C1 (*nonselective*) and C8 (*other organonitrogens*) classes, as shown in Table 4. In contrast, there is no difference between the informative positions in the central one-third of the TMS regions and the remaining exterior regions in the sequences that belong to the C1 (*nonselective*), C2 (*water*), C5 (*organic anions*), C8 (*other organonitrogens*), C9 (*nucleotides*), C10 (*organic heterocyclics*), and C11 (*miscellaneous*) classes; the difference is significant in the sequences that belong to the C3 (*inorganic cations*), C4 (*inorganic anions*), C6 (*organo-oxygens*), and C7 (*amino acids and derivatives*) classes, as presented in Table 5.

---

**Fig 1.**
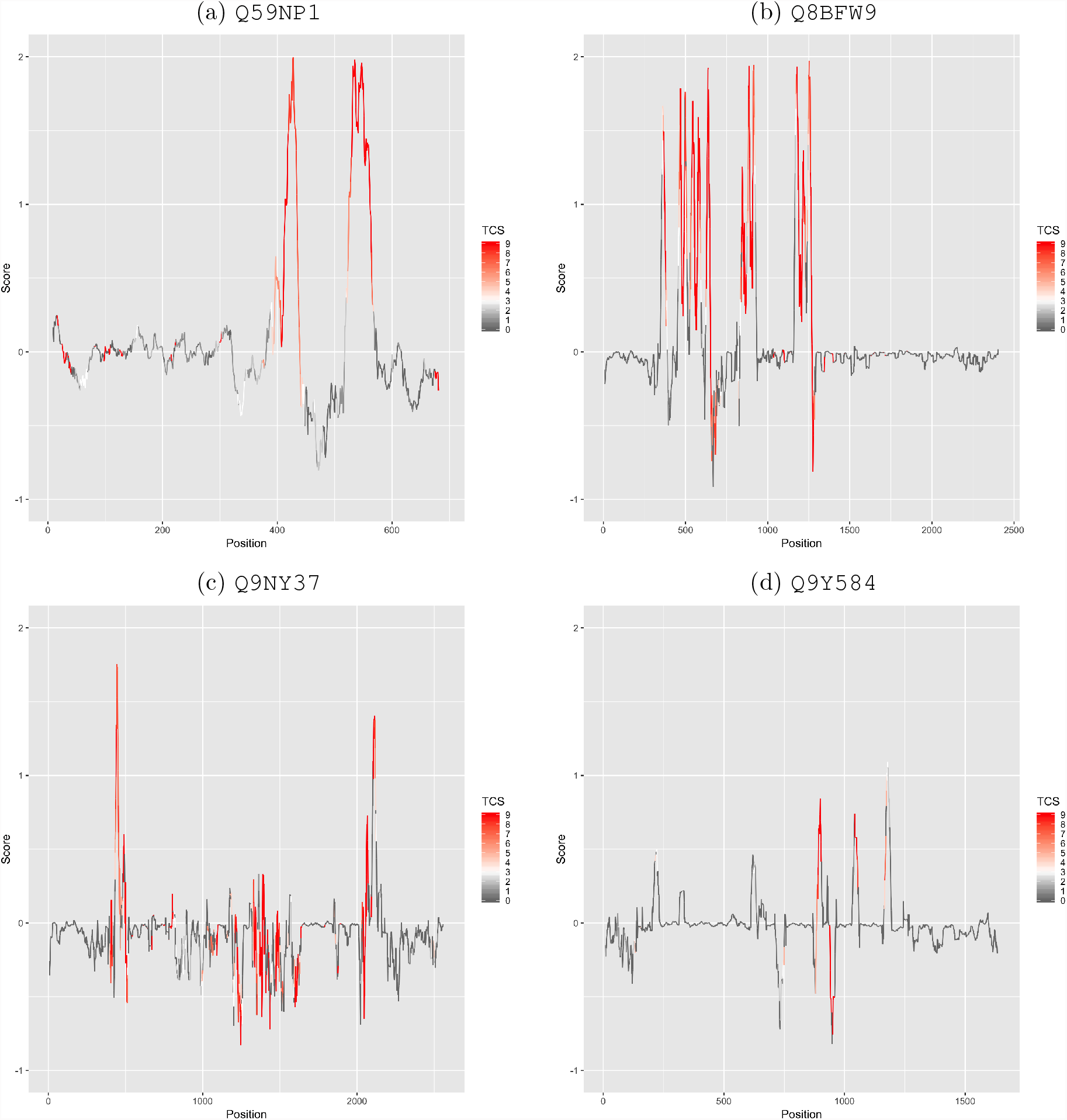
Average Kyte-Doolittle hydropathy of the MSAs with TCSs. The figure indicates that the columns highlighted in red are informative and used by *TooT-SC*. The *TooT-SC* considers a column to be informative if it has a TCS of at least 4 (shades of red) and filters out the other columns (gray and white). In (a), `Q59NP1` contains 251 residues, and the alignment of `Q59NP1` with other homologous sequences has 692 columns; only 151 of them are informative (highlighted in shades of red). In (b), `Q8BFW9` contains 622 residues, and the alignment of `Q8BFW9` with other homologous sequences has 2,414 columns; only 439 of them are informative. In (c), `Q9NY37` contains 505 residues, and the alignment of `Q9NY37` with other homologous sequences has 2,568 columns; only 508 of them are informative. In (d), `Q9Y584` contains 194 residues, and the alignment of `Q9Y584` with other homologous sequences has 1,644 columns; only 79 of them are informative.

**Table 1.** Positional information. This table presents information on the sites retained by the TCS filtering step. For each class of substrates in the dataset, the table presents the average sequence length (**SeqLth**), the average number of TMS regions (**TMS**), and the average total number of residues in the TMS regions (**TMSLth**). It also presents the average of the number of positions retained by the filtering step (**Positions: Num**) and the average of the number as a percentage of the total sequence length (**Positions: %Seq**). It notes the total number of sites that occur in the TMS regions (**TMS: Num**) and the non-TMS regions (**non-TMS: Num**). For the TMS regions, it presents the average number of informative sites that occur in the central one-third of the TMS regions (**TMS: Interior: Num**), and in the remaining exterior regions outside of the central one-third of the TMS regions (**TMS: Exterior: Num**). For the non-TMS regions, it presents the average number of informative sites that occur close to the TMS regions (within 10 positions of the TMS) (**non-TMS: Close: Num**) and the remaining sites far from the TMS regions (**non-TMS: Far: Num**).

| Class ID | SeqLth | TMS | TMSLth | Positions |  | TMS |  |  | Non-TMS |  |  |
| --- | --- | --- | --- | --- | --- | --- | --- | --- | --- | --- | --- |
|  |  |  |  | Num | %Seq | Num | Interior Num | Exterior Num | Num | Close Num | Far Num |
| C1 | 322 | 4 | 81 | 200 | 64.35 | 63 | 22 | 41 | 138 | 35 | 103 |
| C2 | 273 | 6 | 126 | 203 | 74.72 | 121 | 42 | 79 | 82 | 57 | 25 |
| C3 | 681 | 7 | 149 | 387 | 57.23 | 126 | 45 | 81 | 250 | 65 | 185 |
| C4 | 575 | 8 | 168 | 376 | 62.01 | 142 | 50 | 92 | 215 | 73 | 142 |
| C5 | 598 | 10 | 203 | 417 | 70.69 | 179 | 62 | 117 | 233 | 91 | 142 |
| C6 | 461 | 10 | 203 | 325 | 70.45 | 177 | 62 | 115 | 144 | 70 | 74 |
| C7 | 467 | 10 | 206 | 306 | 67.33 | 170 | 59 | 111 | 136 | 83 | 53 |
| C8 | 537 | 4 | 83 | 347 | 39.34 | 70 | 24 | 46 | 133 | 37 | 96 |
| C9 | 403 | 6 | 129 | 282 | 71.25 | 122 | 43 | 79 | 159 | 79 | 80 |
| C10 | 497 | 12 | 241 | 402 | 79.86 | 218 | 76 | 142 | 183 | 95 | 88 |
| C11 | 639 | 7 | 149 | 349 | 47.44 | 110 | 38 | 72 | 164 | 54 | 110 |

**Table 2.** Examples of the informative residue distributions with respect to TMSs and non-TMSs. This table shows the details of individual sequences in Figure 1. The table presents the sequence length (**SeqLth**), the number of TMS regions (**TMS**), and the total number of residues in the TMS regions (**TMSLth**). It also presents the number of informative positions retained by the filtering step (**Positions: Num**) and that number as a percentage of the total sequence length (**Positions: % Seq**). It also denotes the total number of informative sites that occur in the TMS regions (**TMS: Num**), as well as that number as a percentage of the total TMS length (**TMS: % Seq**). In addition, the total number of informative sites that occur in the non-TMS regions (**non-TMS: Num**) are reported, as well as that number as a percentage of the total non-TMS length (**non-TMS: % Seq**).

| UniProt-ID | SeqLth | TMS | TMSLth | Positions |  | TMS |  | non-TMS |  |
| --- | --- | --- | --- | --- | --- | --- | --- | --- | --- |
|  |  |  |  | Num | % Seq | Num | % Seq | Num | % Seq |
| Q59NP1 | 251 | 2 | 42 | 103 | 41.04 | 42 | 100.00 | 61 | 29.19 |
| Q8BFW9 | 622 | 12 | 252 | 386 | 62.06 | 246 | 97.62 | 140 | 37.84 |
| Q9NY37 | 505 | 2 | 42 | 355 | 70.30 | 31 | 73.81 | 324 | 69.98 |
| Q9Y584 | 194 | 3 | 63 | 78 | 40.21 | 32 | 50.79 | 46 | 35.11 |

**Table 3.** Statistical analysis of the informative position rates in the TMS and non-TMS regions. All of the data are reported as the sample mean *±* SD. The locations of the TMS regions are shown as annotated by the `Swiss-Prot` database. There are statistically significant (P-value <0.0001) informative positions in the TMS regions compared to the non-TMS regions in the sequences from all classes except for the *nonselective* class, where the difference is not significant.

| Class ID | TMS | non-TMS | P-value |
| --- | --- | --- | --- |
| C1 | 80.74±23.46 | 58.69±22.43 | 0.0007 |
| C2 | 95.58±9.43 | 57.48±12.14 | $<0.0001$ |
| C3 | 78.31±28.07 | 49.57±22.49 | $<0.0001$ |
| C4 | 79.81±27.38 | 53.94±25.36 | $<0.0001$ |
| C5 | 88.74±20.17 | 60.55±19.79 | $<0.0001$ |
| C6 | 85.35±15.54 | 56.20±16.58 | $<0.0001$ |
| C7 | 81.95±16.90 | 55.28±17.58 | $<0.0001$ |
| C8 | 46.18±44.77 | 34.39±33.03 | $<0.0001$ |
| C9 | 94.67±6.00 | 59.84±6.84 | $<0.0001$ |
| C10 | 90.45±14.48 | 69.15±17.63 | $<0.0001$ |
| C11 | 55.77±37.82 | 41.13±27.80 | $<0.0001$ |

**Table 4.** Statistical analysis of the informative position rates close to TMS regions and far from TMS regions. All of the data are reported as the sample mean *±* SD. For the non-TMS regions, there are statistically significant (P-value <0.0001) informative positions that occur close to the TMS regions (within 10 positions of the TMS) compared to other regions far from TMS regions in the sequences that belong to most classes, except the C1 (*nonselective*) and C8 (*Other organonitrogens*) classes, where the differences are not significant.

| Class ID | Close | Far | P-value |
| --- | --- | --- | --- |
| C1 | 78.24±23.09 | 53.31±26.22 | 0.002 |
| C2 | 76.58±10.97 | 38.59±15.94 | $<0.0001$ |
| C3 | 66.82±26.47 | 43.77±22.95 | $<0.0001$ |
| C4 | 67.26±26.48 | 47.89±26.31 | $<0.0001$ |
| C5 | 78.15±19.54 | 50.94±21.79 | $<0.0001$ |
| C6 | 69.96±14.50 | 45.09±19.63 | $<0.0001$ |
| C7 | 69.18±17.71 | 43.39±20.65 | $<0.0001$ |
| C8 | 38.10±41.33 | 30.53±30.93 | 0.001 |
| C9 | 76.60±06.79 | 49.55±11.43 | $<0.0001$ |
| C10 | 80.52±14.54 | 58.05±23.81 | $<0.0001$ |
| C11 | 49.91±33.30 | 34.75±26.89 | $<0.0001$ |

**Table 5.** Statistical analysis of the informative position rates in the interior and exterior TMS regions. All of the data are reported as the sample mean *±* SD. For the TMS regions, there is no difference between the informative positions in the central one-third of the TMS regions and the remaining exterior regions in the sequences that belong to the C1 (*nonselective*), C2 (*water*), C5 (*organic anions*), C8 (*other organonitrogens*), C9 (*nucleotides*), C10 (*organic heterocyclics*), and C11 (*miscellaneous*) classes. The difference is significant in the sequences that belong to the C3 (*inorganic cations*), C4 (*inorganic anions*), C6 (*organo-oxygens*), and C7 (*amino acids and derivatives*) classes.

| Class ID | Interior | Exterior | P-value |
| --- | --- | --- | --- |
| C1 | 80.66±24.30 | 80.21±23.55 | 0.6485 |
| C2 | 98.44±07.03 | 94.92±10.54 | 0.0003 |
| C3 | 80.92±28.99 | 77.48±28.05 | $<0.0001$ |
| C4 | 81.74±28.33 | 79.10±27.18 | $<0.0001$ |
| C5 | 90.09±19.91 | 88.25±20.49 | 0.0001 |
| C6 | 87.65±17.15 | 84.68±15.50 | $<0.0001$ |
| C7 | 83.93±17.22 | 81.31±16.97 | $<0.0001$ |
| C8 | 47.03±45.76 | 45.86±44.65 | 0.0641 |
| C9 | 97.82±4.81 | 93.32±6.95 | 0.0001 |
| C10 | 92.73±14.89 | 89.75±14.52 | 0.0002 |
| C11 | 56.88±39.33 | 55.45±37.52 | 0.03335 |

